# Non-linear relationships between auditory mismatch responses and the inharmonicity of complex sounds

**DOI:** 10.1101/2025.09.14.676123

**Authors:** Aniela Brzezińska, Bartosz Witkowski, Małgorzata Basińska, Tomasz Domżalski, Krzysztof Basiński

## Abstract

Predictive processing accounts of perception suggest that the brain generates predictions about the incoming stimuli. Precision-weighting, is an important facet of predictive processing theories and has been extensively studied with mismatch responses using electroencephalography. Harmonicity is a feature of sound that is important for auditory perception and previous research has shown that it modulates the brain’s mismatch responses. Since inharmonic sound spectra contain more information (have higher information entropy), inharmonicity has been suggested to be involved in precision weighting. In this study we explored this issue by parametrically modulating the level of inharmonicity applied to synthetic sounds and recording mismatch responses (MMN and P3a) from healthy volunteers (N = 37). Our results show that a sigmoid function models the relationship between inharmonicity and MMN amplitude better than any linear or polynomial function. Furthermore, P3a amplitude has an inverted-U relationship with inharmonicity and peaks at inharmonicity levels just below the threshold for pitch discrimination. These results are consistent with the hypothesis that inharmonicity impairs F0 extraction above a certain threshold and does not serve as an index of precision in the auditory system.

## Introduction

If a complex tone consists of frequencies that are integer multiples of one fundamental frequency (F0), it is considered harmonic. Conversely, complex tones that include non-integer multiples of one frequency are considered inharmonic. Harmonicity is important in different domains of auditory perception. Many ecologically salient sounds have harmonic spectra - these include human speech, animal vocalizations, or sounds of musical instruments. Inharmonic sounds are harder to detect in noise, suggesting a role in auditory scene analysis^1–3^. Pitch, the perceptual attribute that allows sounds to be segregated on the scale from low to high, largely depends on harmonicity ^4–8^. Some pitch discrimination tasks are severely impaired for inharmonic sounds ^9^. Finally, harmonic sounds may be more efficiently encoded in memory because their spectra can be easily compressed by extracting and representing the F0 10. F0 extraction is a process where the auditory system computes the most likely F0 given a set of simultaneously perceived frequencies (allowing for perception of F0 even when no acoustic energy is present at that frequency) ^5^.

Predictive processing (PP) accounts of perception suggest that the brain is using an internal, generative model to form predictions about the incoming stimuli ^11–18^. Whenever these predictions are violated, a prediction error response is elicited that reflects the difference between the perceived stimulus and the prediction. Prediction errors in PP are learning signals and serve to change the contents of the generative model. However, the environment is extremely rich in information, and much of that information is irrelevant. This presents a challenge to sensory systems that need to filter out relevant information and ignore noise. To solve this problem, prediction errors are thought to be weighted by the self-estimated uncertainty of perceptual inference - *precision* ^13,19,20^. This hypothetical mechanism allows for down-weighting of prediction errors arising from noisy, incomplete or ambiguous stimuli, thereby not allowing these errors to influence future predictions. A popular PP formulation suggests that precision is proportional to the information content of the stimulus (stimuli containing more information are more difficult to predict) ^13,18,21^. Precision weighting is an important facet of PP and has been extensively studied in recent years ^22–28^. In the human auditory system, this process can be studied by employing oddball paradigms and recording event-related potentials (ERPs) using electroencephalography (EEG) ^29,30^. The mismatch negativity (MMN) is a widely studied ERP component that is established as an electrophysiological trace of precision-weighted prediction errors in the cortical auditory system ^21,31–36^. MMN is often followed by P3a, a positive ERP component associated with an automatic attention shift to the perceived mismatch ^37^.

The spectrum of a harmonic sound can be reliably represented by the harmonic series starting at the F0. Inharmonic sounds have spectra that include mathematically unrelated frequencies. Thus, in information-theoretic terms^38^, inharmonic sounds carry more information (they have higher information entropy) than harmonic sounds. Any prediction errors elicited by inharmonic sounds should therefore be down-weighted in precision, due to their increased information content. This, in conjunction with psychophysical results suggesting impaired perception of inharmonic sounds and impaired retention of inharmonic sounds in memory ^2,9,10^, has led to the hypothesis that harmonicity can potentially serve as an index of precision in the auditory system ^29,39^. A recent study explored this idea empirically by exposing participants to harmonic and inharmonic sounds in a roving oddball paradigm ^40^. Using synthetic stimuli, the authors induced inharmonicity to the harmonic complex tone by introducing a random jitter to all frequencies above the F0. Surprisingly, inharmonic sounds with a constant jittering pattern throughout the sequence generated similar MMN and larger P3a responses than harmonic sounds. However, when the jittering pattern was randomized for each consecutive sound, both MMN and P3a responses were not detectable. This finding is consistent with precision weighting, as inharmonicity leads to a decrease in mismatch responses, at least for sequentially changing sounds. However, since the study compared harmonic to heavily jittered inharmonic sounds, it is unclear if the effect is caused by precision weighting (an increase in the uncertainty of predictions) or a fluctuating perception of the standard’s pitch caused by the impaired F0 extraction. For a sufficiently jittered inharmonic sound, F0 extraction might be impossible because the spectrum cannot be approximated by any single harmonic series ^40^.

In the current study, we aimed to explore the relationship between inharmonicity and precision-weighting further. We prepared synthetic sound sequences with randomly jittered frequencies and systematically manipulated the rate of this jitter. The increase in the level of sound inharmonicity directly corresponds to an increase in information entropy of the acoustic signals. We recorded MMN and P3a mismatch responses to pitch deviants in these sequences using EEG. We investigated two competing hypotheses. The primary hypothesis states that precision-weighted prediction errors to sounds that increase in inharmonicity theoretically should manifest in a linear (or transformed-to-linear) decrease of MMN amplitudes ^13,31,33,34^. This is because the gradual introduction of inharmonicity increases the information content of sounds and, all else being equal, makes any prediction more uncertain (less precise). An alternative hypothesis assumes that the auditory system employs F0 extraction to detect deviants. In this case, we expect the MMN amplitudes to remain similar for small jitter rates, as F0 extraction would not be impaired by low-to-moderate levels of inharmonicity. Once a threshold value is reached, we expect mismatch responses to drop abruptly to near-zero levels (undetectable MMN) for larger jitters, presenting as a sigmoid relationship. This is because if the F0 extraction is impaired, the perception of regularity itself becomes unstable and any incoming deviance would not elicit a mismatch response.

Our results show that a sigmoid function modeled the relationship between inharmonicity and MMN amplitude data better than any linear or polynomial function. This is consistent with the hypothesis that inharmonicity impairs F0 extraction above a certain threshold and may not serve as an index of precision in the auditory system. These findings are further supported by behavioral data, showing that pitch discrimination was impaired at a similar inharmonicity threshold. Finally, we also observed a slight peak in the P3a responses around the inharmonicity threshold which can reflect the higher computational demand associated with detecting pitch deviants at the threshold level.

## Results

### Procedure outline

In order to measure the MMN and P3a mismatch responses, we performed an EEG experiment using a version of the roving oddball paradigm ^34,41^. This paradigm involves presenting a sequence of repeating sounds at a specific F0, followed by another sequence at a different, randomly chosen F0, followed by another sequence and so on. The first sound after the F0 shift generates a mismatch response, but it becomes the new standard after several repetitions (Figure 1A). The roving paradigm ensures that standards and deviants have, on average, the same acoustical properties, with mismatch induced by the context of the sequence.

**Figure 1.**
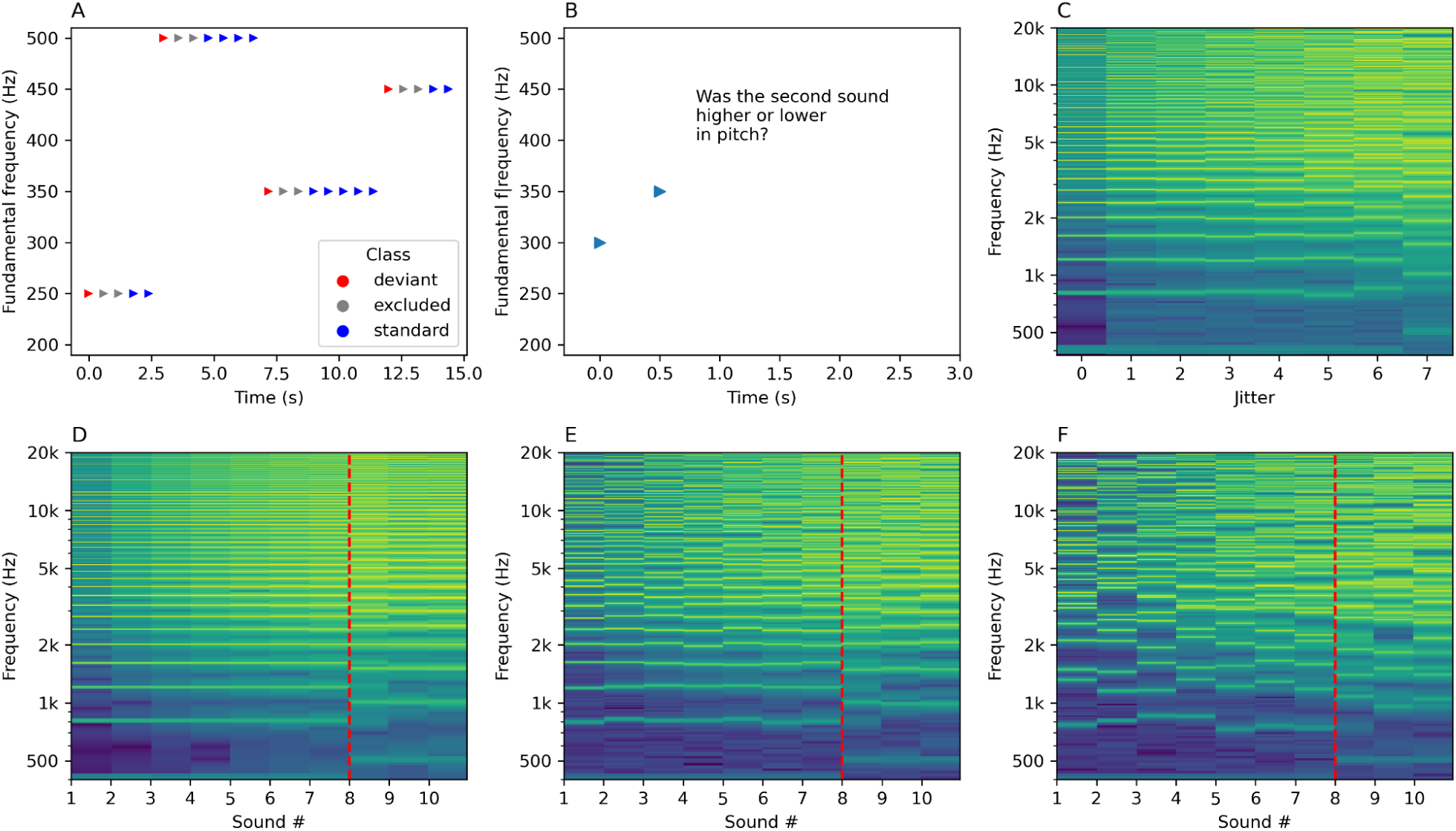
Experimental paradigms. Panel A illustrates the roving oddball paradigm. The first sound after a switch in fundamental frequency is treated as a deviant (in red). After a few repetitions, the deviant becomes a new standard (blue). Two sounds immediately after the deviant are excluded from the analysis (grey). Panel B illustrates the behavioral task where a pair of sounds is presented and the participant is required to determine if the second sound was higher or lower in pitch than the first. Panels C - F show spectral decompositions of sounds as a function of time/jitter and frequency. Brighter colors indicate more acoustic energy at a particular frequency. Panel C presents spectrograms of example sounds used in both experiments. The increasing jitter (x-axis) introduces increasing spectral complexity to frequencies above the F0 (in this example 400 Hz). Panels D-F display the spectral structure of example sound sequences used in the roving oddball paradigm for jitter condition *j0* (Panel D), *j4* (Panel E) or *j7* (Panel F). The x-axis represents time, and the y-axis represents frequency in Hz. For clarity of presentation, the silence between the sounds was removed, so the time axis does not reflect the actual IOI (600 ms). Red vertical lines mark the point where the F0 changes, introducing a pitch deviance. For higher jitter rates, the spectral structure of the sounds becomes increasingly irregular.

To systematically test the mismatch responses to progressively higher amounts of inharmonicity, we manipulated the level of inharmonicity between the experimental conditions. In the condition *j0* (“jitter 0”) we used harmonic complex tones. For conditions *j1 - j10*, we applied random jittering to all frequencies above the F0, for each sound in the sequence (see Figure 1C-F for examples). The jittered frequencies F_n_’ were calculated using the following formula:

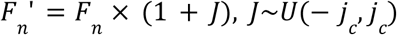

where *F_n_* is the frequency of the n-th harmonic and *J* is a continuous uniformly distributed random variable ranging from *-j_c_* to *j_c_*, with *j_c_* being the condition-specific jitter rate. The upper limit of the jitter rate, i.e. *j10* = 0.5 is the highest possible jitter value that does not lead to a reordering of frequency components. Pilot data indicated that if we used jitter rates that increase linearly from zero to 0.5, pitch discrimination will be at chance level for the participants in the *j1* condition. Thus, we decided to choose jitter rates that progress geometrically, increasing the resolution of manipulations for the lower jitter levels. In order to reduce recording times, we did not test conditions *j8 - j10* with EEG, as pilot data as well as previous research ^40^ indicated that they likely not elicit a detectable MMN. We also applied a stationary high-pass filter to all sounds in order to attenuate the low frequencies and focus the auditory processing on upper harmonics, unresolved by the cochlea (see Method for details of this procedure).

### Mismatch negativity analysis

In the EEG experiment, a typical pattern of MMN and P3a responses was found in the harmonic condition (a fronto-central negativity in the 100-200 ms latency range followed by a later fronto-central positivity). These responses diminished with increasing jitter (Figure 2, Figure 3). Mean MMN amplitudes differed significantly from zero for all conditions (means and t-test results are reported in Supplementary Table 1). One-way repeated measures ANOVA showed that harmonicity affected MMN amplitudes (F(7, 238) = 9.12, p_Greenhouse–Geisser_ < 0.001, Supplementary Table 2). To explore pairwise differences between jitter conditions, post-hoc comparisons were performed. MMN amplitude differed significantly between lower and higher levels of inharmonicity. Specifically, jitter levels *j0*, *j1*, *j2*, and *j3* differed significantly from *j5*, *j6*, and *j7*, (with p-values ranging from p = .0044 to p < .0001; see Supplementary Table 3), indicating a significant reduction in MMN amplitude at higher jitter levels. However, pairwise comparisons between low jitter levels (*j0–j4*) were not statistically significant (all p-values > .05). Similarly, pairwise comparisons between all higher jitter levels (*j5–j7*) were also not statistically significant (all p-values > .05). These results suggest that the most pronounced changes in MMN occurred between the *j4* and *j5*, rather than incrementally between all adjacent levels (Figure 3, Supplementary Table 3). This pattern of results is consistent with a model suggesting an abrupt decrease in the strength of mismatch responses at the threshold value, rather than a gradual change (as in precision weighting).

**Figure 2.**
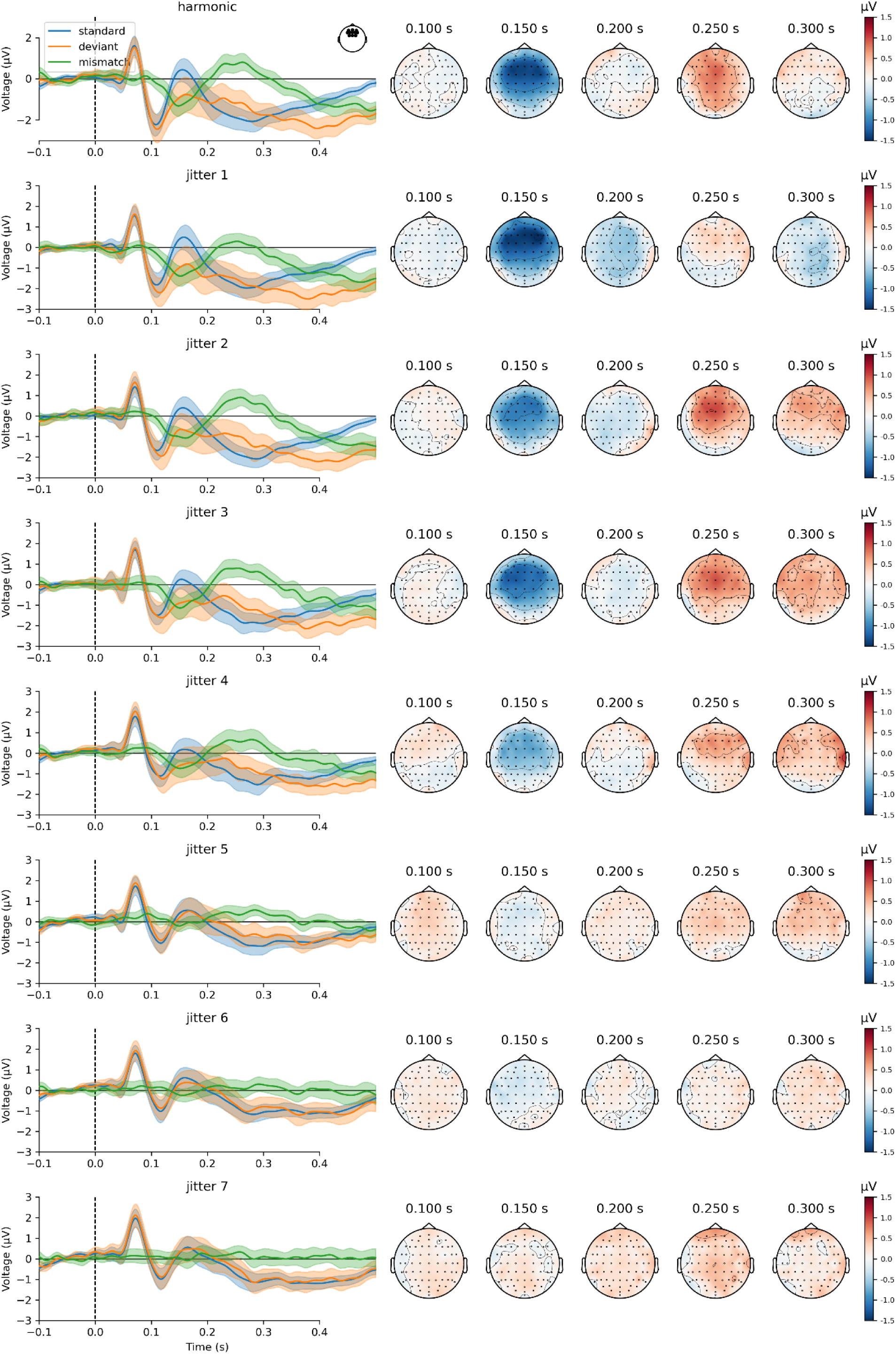
Grand-averaged event-related potentials and topographies of whole-brain activity. Each row represents an experimental condition, starting with harmonic (first row) and followed by seven inharmonic conditions. Event-related potential traces (left) show grand-average responses to standards (blue), deviants (orange), and the difference waves (green) for fronto-central channels (Fz, F1, F2, AF3, AF4, AFz, F3, F4, FC1, FC2, FCz). Color-shaded areas represent 95% confidence intervals of the mean. Scalp topographies (right) present signal strength at chosen latencies.

**Figure 3.**
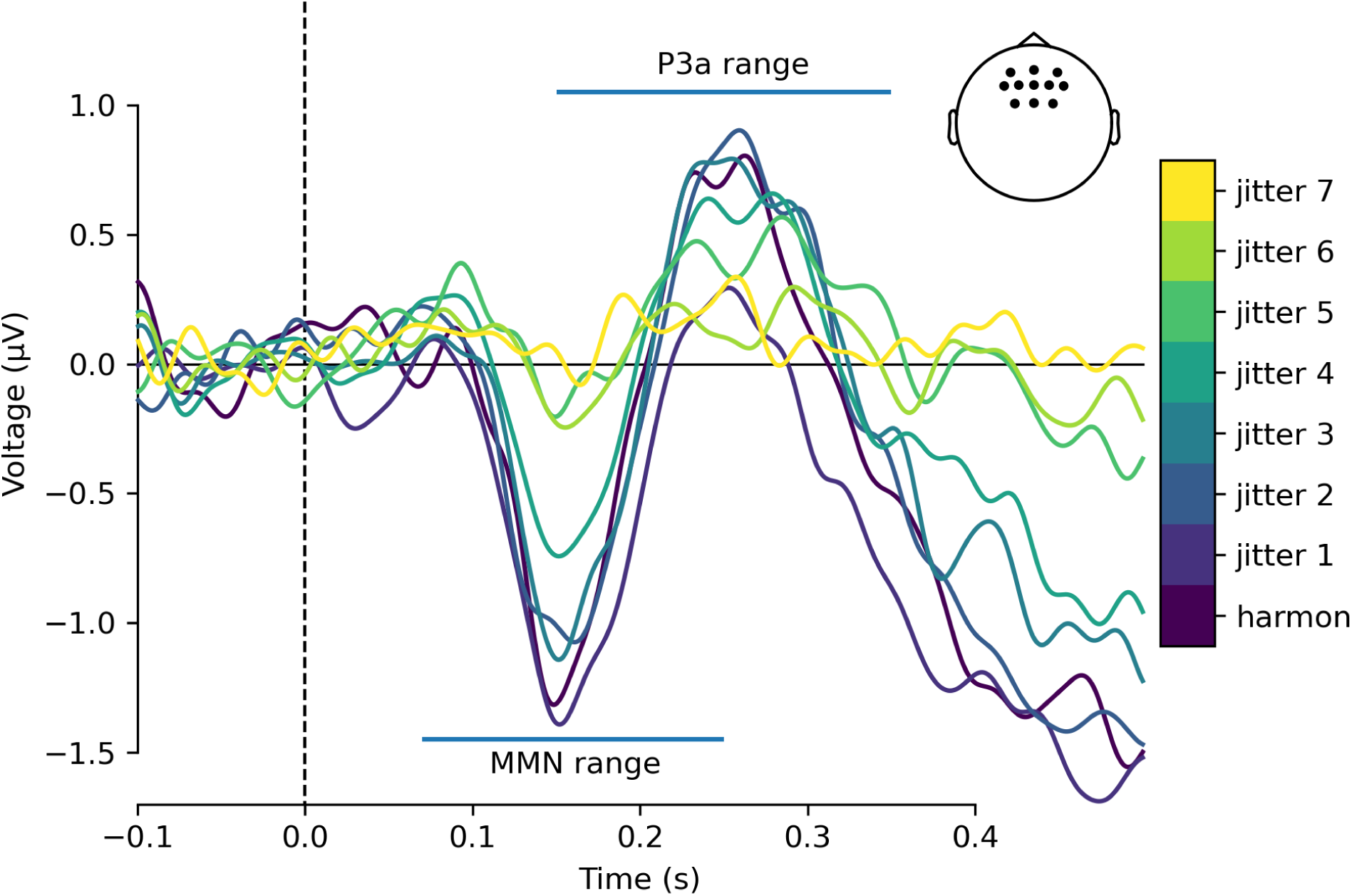
Grand-average mismatch wave differences between experimental conditions. Each trace is a difference wave between deviant and standard response in a given jitter condition. Traces present averaged activity in the fronto-central channels (Fz, F1, F2, AF3, AF4, AFz, F3, F4, FC1, FC2, FCz). Confidence intervals were not shown for visual clarity but can be seen in Figure 2. Horizontal lines indicate MMN and P3a peak latency windows.

To further explore how MMN amplitude depends on sound harmonicity, we used linear, polynomial and non-linear mixed effects models with differential entropy of jitter distribution as an independent variable. Since the jitters were drawn from a uniform distribution, with support ranging from -*j_c_* to *j_c_*, where *j_c_* was condition specific jitter rate, differential entropy *I* was expressed as:

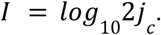

This parametrization allowed us to quantify the amount of information (or uncertainty) contained in the sounds while, at the same time, effectively linearizing the jitter rates and avoiding possible issues with fitting models to very small jitter rates. Of note, differential entropy is not an exact continuous analogue of discrete entropy and does not share all its properties. Differential entropy can assume negative values, as was the case in our experiment, and zero differential entropy does not indicate certainty. For an outcome that is certain, differential entropy is not defined so in the further analyses we omit the harmonic condition with zero jitter rate.

We conjectured that the relation between inharmonicity and MMN amplitude is sigmoidal rather than linear and can be modeled as the following function:

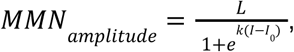

where *I* is differential entropy of jitter sampling distribution, *I*_0_ is the *I* value at the inflection points, *k* determines the slope at the inflection point and *L* is the left asymptote of the MMN amplitude (right asymptote is at 0, reflecting the disappearance of MMN for highly inharmonic sounds). Likelihood ratio test showed that the sigmoid mixed effects model with random left asymptote showed significantly better fit than the null model (χ^2^(2) = 75.48, p<0.001). Linear model with random intercept also showed significantly better fit to the data than the null model (χ^2^(1) = 46.90, p < 0.001). Adding higher degree parameters to the model did not significantly improve model fit (see Supplementary Table 4 for details). At the same time, both the Akaike and Bayesian information criteria were markedly better for the sigmoid model (AIC = 664.6, BIC = 682.2) than for the linear model (AIC = 691.2, BIC = 705.2), which suggests that the relationship between inharmonicity and MMN amplitude is indeed better modelled by a sigmoid than a linear or polynomial function. This corroborates the results of the post-hoc contrast analysis. Since the linear and sigmoid models were not nested, we refrain from reporting the *p* value for the likelihood ratio test for the comparison of these two models, as the distribution of the likelihood ratio for non-nested mixed models is not known and various distributions are possible ^42^.

The I_0_ parameter of the fitted curve was −1.18, suggesting that the MMN decays rapidly for jitter rate around 0.033, that is between *j4* and *j5*, convergent with the results of the contrasts analysis. The fitted curve is presented in Figure 4C. The details of both sigmoidal and linear models plots are reported in the Supplementary Table 5.

**Figure 4.**
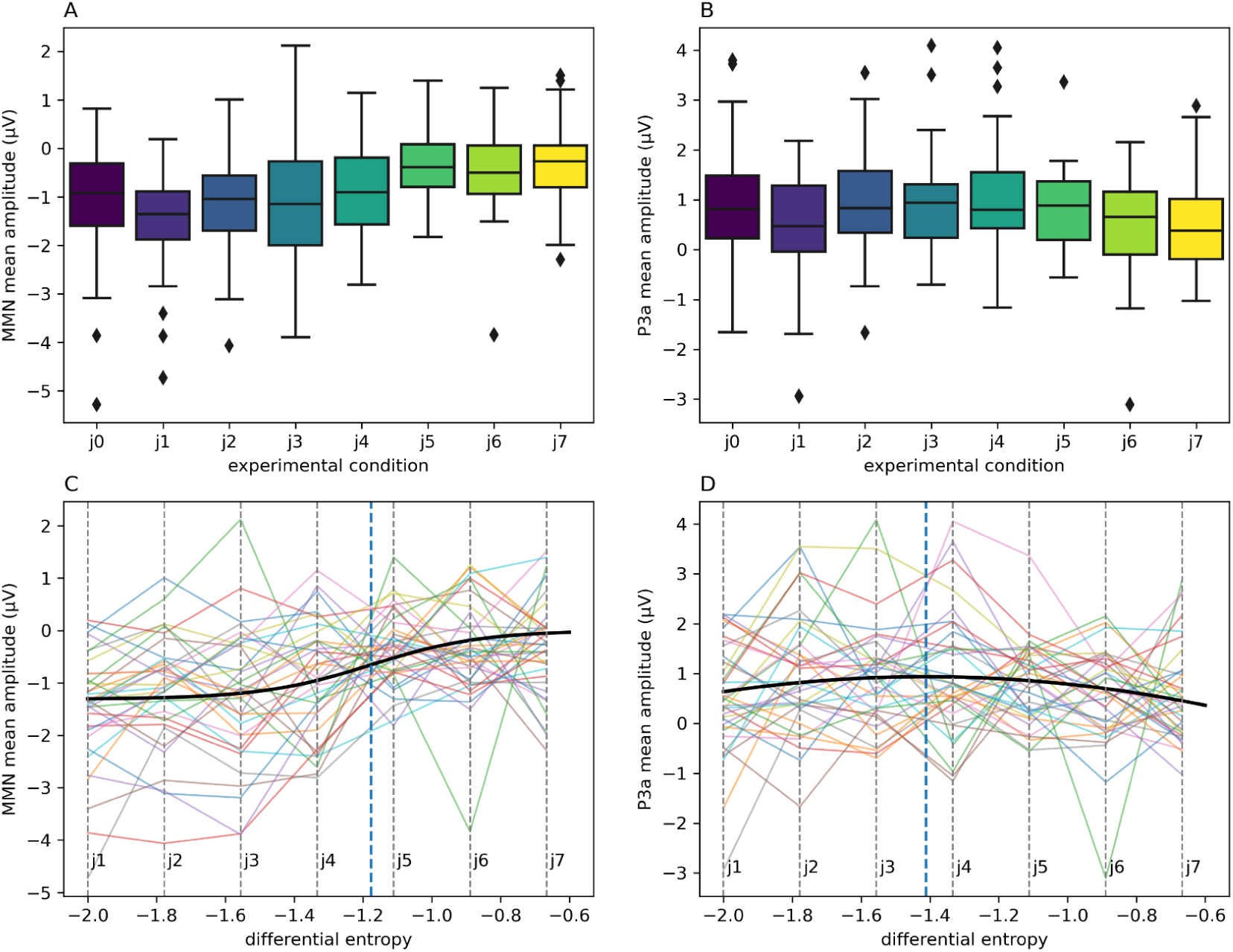
Peak measures differences between conditions. Panel A shows MMN mean amplitude, Panel B shows P3a mean amplitude. Box plots show quartile distributions of obtained results, whiskers indicate minima/maxima while diamonds indicate outlier observations. Panel C presents a sigmoid curve (black line) representing a function modelling the relationship between the differential entropy of jitter factor distribution and MMN amplitude, fitted on the experimental data. Panel D presents a quadratic curve fitting the P3a amplitude data. Vertical dashed lines in Panels C and D correspond to experimental conditions j1-j7. The blue dashed line represents the sigmoid inflection point (Panel C) or the quadratic function maximum (Panel D). Colored lines connect observations from the same participant.

MMN peak latency was not significantly related to sound harmonicity, both in ANOVA (F(7, 238) = 0.82, p_Greenhouse–Geisser_ = .538, see also Supplementary Table 2) as well as in linear, quadratic and cubic mixed models (all p-values > .05, see also Supplementary Tables 4 and 5).

### P3a analysis

P3 amplitudes (Figure 4D) differed significantly from zero for all harmonicity conditions (Supplementary Table 1). ANOVA results did not indicate any significant effect of condition (F(7, 238) = 1.78, p_Greenhouse–Geisser_ = 0.113, Supplementary Table 2). However, we observed a significant relationship between the differential entropy of the jitter rate distribution and P3 amplitude in the mixed models analysis. The quadratic model had a significantly better fit than both null (χ^2^(2) = 6.93, p = 0.031) and linear models (χ^2^(1) = 5.95, p = 0.015). The quadratic function had a shape of an inverted parabola with a predicted maximum at entropy = −1.41, corresponding to jitter conditions between *j3* and *j4.* These results suggest that P3a amplitudes form an “inverted-U” relationship with differential entropy of the stimulus, peaking at moderate entropy values while remaining low for both low and high entropy values.

P3 latency was not significantly related to sound harmonicity, both in the ANOVA analysis (F(7, 238) = 1.22, p_Greenhouse–Geisser_ = 0.299,Supplementary Table 2) and in linear and polynomial mixed models (all p-values > .05). Model comparison results and parameters of the fitted models are reported in ST4 and ST5.

### Behavioral results

In the behavioral experiment (Figure 1B, Supplementary Figure 1), participants were required to perform a two-alternative forced choice pitch discrimination task. For a sequence of two sounds, the participants were to determine if the second sound is higher or lower in pitch than the first sound. We employed an adaptive staircase procedure (3-up-1-down) that modulated the amount of inharmonicity applied to the sounds, starting randomly from either *j0* (harmonic) or *j10* (inharmonic with maximal jittering). This procedure halted after 6 reversals and was performed 15 times for each participant. We took the final reversal jitter levels as indicative of 79.37% discrimination threshold ^43^ and averaged them within participants. The mean jitter value for discrimination threshold was M = 2.84 (SD = 2.32), indicating that the participants had trouble discriminating pitch for jitters from *j3* upwards.

## Discussion

The results of this study show that the relationship between the level of inharmonicity and MMN amplitude can be best modelled with a sigmoid function. This is additionally supported by ANOVA and post-hoc contrast results, showing that MMN amplitude is similar for harmonic and low-inharmonic sounds. At the same time, MMN amplitudes are significantly smaller for highly inharmonic sounds. The sigmoid model parameters and post-hoc comparisons place the threshold above which MMN amplitude suddenly decreases between jitters *j4* and *j5*. Evidence from the behavioral experiment suggests that pitch detection deteriorates for jitters above *j3*. Overall, the presented results are consistent with the hypothesis that inharmonicity does not exert influence on auditory deviance detection until a specific threshold, where it begins to impair F0 extraction. At the same time however, our results cannot serve as definite evidence against theories that suggest that increased information entropy leads to down-weighting of prediction errors by precision. A linear model of the relationship between MMN amplitude and inharmonicity also fit the data better than a null model (albeit the fit for the sigmoid model was better). The interpretation of these findings is also limited by methodological issues, as discussed below.

It is possible that precision-weighting processes are not reflected in the MMN amplitudes or that precision-induced changes are so minute that they are not observable in noisy EEG data. While this is certainly a possibility, recent simulation work suggests that precision weighting should be apparent not only in amplitudes, but also in the latencies of the putative event-related potentials ^44^. We do not find this to be the case in our results, as latencies of both MMN and P3a were not modulated by the level of inharmonicity. However, one needs to remember that different models of how precision-weighting might be implemented in the brain exist in the literature (i.e. gain modulation, variability modulation, synaptic modulation, ^45^ see also ^11^ for review). More simulation work is required to provide concrete predictions about how these different precision-weighting mechanisms might manifest in scalp ERPs.

While our results suggest that harmonicity is not an index for precision-weighting, they do not contradict the models of predictive processing that implement precision-weighting. Rather, they suggest a process where pitch information is extracted from harmonic and mildly inharmonic sounds, but the auditory system struggles to perform this task for higher inharmonicity values. Further, the confidence (precision) of any pitch related predictions does not seem to influence the amplitude or latency of MMN and P3a components. This possibly results from the fact that F0 extraction (and the associated prediction) might be performed earlier in the auditory processing hierarchy.

We observed a peak in the P3a amplitudes between jitters *j3* and *j4*. This peak can reflect a more salient attentional shift ^37,46^, perhaps caused by the higher computational demand associated with detecting pitch deviants near the threshold level. Interestingly, this peak occurred for slightly lower inharmonicity levels than the MMN sigmoid threshold, indicating that this attentional shift is the highest for near-threshold inharmonicity levels. For lower inharmonicity levels, deviant detection is computationally easier and does not capture attention as much. Conversely, for high inharmonicity levels the auditory system does not detect deviants at all (as evidenced by MMN results), therefore diminishing the P3a responses. Deviance detection may also explain the presence of later negativity (post-350 ms) in the low-jitter conditions (Figure 3). Because of the nature of the roving paradigm, sounds immediately following the deviant “carry over” some of the deviant response. For this reason, we chose to exclude the second and third sound after each deviant from the analysis. The divergence disappears for higher jitter conditions (j5-j7), where there is no deviance detection and consequently no prediction error, further supporting this interpretation.

The present study is limited in several aspects. Although the sigmoidal model fitted the data better than the linear model, the result should be interpreted with caution. As can be seen from residual plots (Supplemental Figure 2 and 3), both linear and sigmoid models show signs of heteroschedasticity. Furthermore, the geometric progression of jitter rates used for manipulation might have biased the model comparisons. To mitigate these problems, future work can utilize linear jitter rate parametrization that focuses on the immediate vicinity of the inflection points established here. Finally, while a sigmoid relationship is consistent with our alternative hypothesis (suggesting an impairment in F0 extraction), there are surely many different possible neural mechanisms that could produce a similar pattern of results.

It is important to remember that the results of the present study were obtained from a passive paradigm. This likely contributed to the overall small ERP amplitudes ^47^. With an active task (i.e., responding to pitch deviants), a P3b response might have also been elicited, exerting influence on the P3a. Furthermore, the computations performed by the auditory system might be different in an active task in comparison to a passive one, which could lead to different patterns of mismatch responses (particularly in the P300 complex). Different computations might also underpin different perceptual strategies for detecting deviants, consistent with substantial inter-individual differences observed in the present study in both neuroimaging and behavioral results. Of note, our P3a analysis is exploratory in nature, as we did not formulate any a’priori hypotheses about the nature of this component’s relationship with inharmonicity. Future research could address these issues by utilizing active paradigms on larger sample sizes to verify hypotheses, perhaps facilitated by gamification to counter participant fatigue and support motivation.

We acknowledge that the behavioral results presented in this study are limited. The aim of the behavioral experiment was to gain insight into the detectability of pitch deviants in order to better interpret the EEG results. As such, the behavioral experiment was not designed to provide thorough psychometric testing of pitch discrimination under inharmonicity. Rather, our aim was to verify if the pitch deviants presented in the passive EEG experiment could be detected if participant’s attention was afforded. Specifically, we have made a decision to retain a constant 50 Hz difference between sounds for each pair in the range of 200 - 500 Hz. We recognize that this decision resulted in different perceived pitch intervals for different starting frequencies, with a bias towards smaller perceived intervals for higher pitches. However, 50 Hz was also the minimal F0 change in the roving paradigm and we wanted to make sure that the behavioral results are, first and foremost, interpretable in the context of the EEG results. In that sense, if participants can perform pitch discrimination for sounds differing 50 Hz in F0, they should also be able to detect deviants in a roving paradigm where differences are the same or larger. A full examination of the effects of inharmonicity on pitch discrimination would involve testing different intervals and different inharmonicity levels independently and is outside the scope of this study.

In sum, the results of the present study are consistent with the hypothesis that inharmonicity impairs F0 extraction above a certain threshold. This is evidenced by the sigmoid relationship between inharmonicity and MMN amplitude. The results are further complemented by the P3aamplitude peaking near the MMN amplitude function inflection point. This may be due to the early auditory system collapsing the harmonic information into an F0-like representation or a lack of sensitivity of scalp EEG to precision signals. However, our results do not speak directly to the neural mechanisms of F0 extraction nor rule out the presence of precision-weighting mechanisms in the auditory system. This study suggests a descriptive model of the relationship between inharmonicity and the auditory mismatch responses and, as such, does not provide any mechanistic explanation for the underlying neural processes. Further research using computational modelling, animal studies using invasive methods, as well as more human neuroimaging data is required to disentangle the mechanisms involved.

## Methods

### Participants

37 people signed up for the study. Inclusion criteria were: no reported neurological or psychiatric illness, age between 18 and 45 years old, normal hearing, and no use of medication that affects the central nervous system 24 hours prior to testing (e.g., opioids, pain medications). The participants were subjected to a tonal audiometry test before the experiment to screen for hearing disorders. Two of the participants did not meet the inclusion criteria, with hearing thresholds below the −20 dB HL (hearing level) benchmark. The remaining 35 did not have problems with hearing and took part in the study. Two participants had to perform the experiment in two sessions, due to tension headaches from the EEG and headphones placed on their heads. The participants were aged between 18 and 37 years (M = 23.7, SD = 3.17). 27 (77%) of them were women, 8 (23%) were men. Participants received monetary compensation for their involvement in the study.

Before data collection, we performed a power analysis to guide sample size decisions. We have simulated MMN peak amplitude data assuming a worst-case scenario where all conditions did not differ except one that was, on average, 1 µV larger than the rest (a difference of this magnitude was observed between conditions in ^40^). We ran this simulation 10 000 times for each sample size in the range between 20 and 40. For each simulated dataset, we fit a mixed-models ANOVA with condition as a fixed effect and participant as a random effect. The results of these simulations revealed that in order to achieve statistical power of 0.8 or more, at least N = 34 participants were required.

### Procedure

The protocol for this study was approved by the Scientific Research Bioethics Committee of the Medical University of Gdańsk (reference number KB/489/2024) and followed the Declaration of Helsinki. Before the procedure, all participants provided written informed consent.

Each participant took part in the following procedures: (1) tonal audiometry - in order to check that the participant has healthy hearing, (2) the behavioral experiment and (3) the EEG experiment. Experiments were performed on a computer running Psychopy v.2023.2.3 (for behavioral) or v.2024.2.1 (for EEG). The sound stimuli were presented binaurally with Beyerdynamic DT 770 Pro headphones and an RME UCX II audio interface.

In the behavioral experiment, participants performed a two-alternative forced choice pitch discrimination task. They were listening to pairs of sounds (inter-onset interval, IOI = 500 ms) and were asked to determine whether the second sound was higher or lower in pitch than the first sound. The level of inharmonicity was manipulated with a 3-up 1-down staircase procedure. The sounds’ F0s were randomly chosen from a range of 200 Hz - 500 Hz in increments of 10 Hz. The difference in frequency between sounds was always set at 50 Hz, the minimal difference in F0 present in the EEG paradigm (see below). Each staircase block ended after 6 reversals and each participant completed 15 blocks. Before the main experiment, a familiarization trial was run with a similar task and using harmonic sounds. The participant proceeded to the main task after three consecutive correct responses in the familiarization trial. Participants responded by pressing a button on the keyboard.

Scalp EEG potentials were collected with a 64-channel active system (BioSemi B.V.) with two additional EOG electrodes. The EEG experiment consisted of 16 blocks, two per inharmonicity condition, each containing roving sequences of 600 sounds played with an IOI of 600 ms. The first tone of a new stimulus train served as the deviant stimulus and a source of a mismatch response in the context of the prior stimulus train. After a few repetitions, the deviant tone establishes itself as the new standard until the next F0 change happens. The number of stimuli in a given train varied pseudo-randomly from 4 to 7. Each block lasted about 6 minutes and was repeated twice. The F0s were chosen from the set of frequencies from 200 Hz to 500 Hz, in 50 Hz increments. The participants were asked to refrain from movement during sound stimulation but could take self-paced breaks between the blocks. During EEG recording, the participants watched a movie of their choice, without sound. The entire procedure, including behavioral testing, lasted for about 2 hours.

### Stimuli

In both experiments, stimuli consisted of synthetic complex tones generated with custom Python code, based on similar procedures used in previous studies ^9,10,40^. Each tone was synthesized by adding sine waves at frequencies of F0 and its harmonic frequencies up to the Nyquist limit (24 kHz), in random phase. For inharmonic sounds, random jitter *J* was applied to each frequency above the F_0_. The jittered frequencies F_n_’ were calculated using the formula given in the Results section. The exact jitter values were as follows: *j0* = 0, *j1* = 0.005, *j2* = 0.008, *j3* = 0.014, *j4* = 0.023, *j5* = 0.039, *j6* = 0.065, *j7* = 0.108, *j8* = 0.180, *j9* = 0.300, *j10* = 0.500. Jitters spanning from *j0* to *j7* were used in the EEG experiment. Rejection sampling was used in cases where two neighbouring frequencies differed by less than 30 Hz in order to avoid beating artifacts. For every sound, a fourth-order Butterworth high-pass filter was applied at the cutoff frequency of 3500 Hz, corresponding to the 10th harmonic of the average F0 used throughout both experiments (350Hz). This filter provided attenuation of the F0 in order to minimize the variation at the spectral edge of the fundamental and to minimize the changes to the overall spectrum during F0 transitions (see ^9^ for further discussion of the rationale behind this approach). Importantly, this filter does not remove all frequency content below the cutoff. Rather, it provides a gradual attenuation of low frequencies (24dB/octave), producing a brighter, somewhat buzzing sound but still with a clear, discernible pitch in the harmonic condition. While we acknowledge that the filtering might have resulted in an overall weaker pitch sensation (due to lesser emphasis on resolved harmonics), the same manipulation was applied to all sounds and therefore does not bias the comparisons between conditions. Sounds had a length of 70 ms and were amplitude modulated with 5 ms onset and offset ramps (half-Hanning windows). All sounds were pre-rendered as 16-bit wav files with a sampling rate of 48 kHz and loudness-normalized to −12 LUFS using *pyloudnorm*. For the behavioral experiment, 100 sounds with different jitter patterns were created for each possible F0 and each jitter rate, for a total of 35,000 sounds. For the EEG experiment, 1000 sounds with different jitter patterns were created for each possible F0 and jitter pattern, for a total of 56,000 sounds.

### Data processing and statistical analysis

The ERP data were collected using a 64-channel BioSemi ActiveTwo system (BioSemi, Amsterdam, Netherlands) at a sampling rate of 2048 Hz. All data was decimated by a factor of two before preprocessing and subsequent analyses were performed with a sample rate of 1024 Hz. The oculomotor activity was recorded using two additional EOG electrodes. Raw EEG data underwent high-pass (0.2 Hz), low-pass (30 Hz), and notch (50 Hz) filtering to remove the AC component. We included the notch filter because any low-pass filter will have a pass-band above the cutoff frequency that gradually tapers to zero and, in extreme cases, might include some of the line noise. At the same time, this filtering should not introduce substantial artifacts and bias the results. The data were then segmented into epochs ranging from −100 ms to 500 ms and subsequently processed using the Python library Autoreject (v0.4.24) ^48^ for automatic detection and rejection of faulty channels and epochs. This machine learning algorithm estimates optimal peak-to-peak signal thresholds for each epoch and each EEG sensor. The thresholding leads to epoch-wise interpolation of bad channels or (in case of very noisy signals) epoch rejection. Next, in order to remove oculomotor artifacts, an independent component analysis (ICA) was performed. ICA components were identified for removal through an automated procedure based on the EOG signals; however, all selections were later manually verified. The ICA solution was applied to the data before automatic artifact rejection. Then, the autoreject algorithm was run once again, as recommended by Jas et al. ^48^. In the end, 13 400 out of 321 389 of epochs had to be rejected (4.2%). The remaining epochs were re-referenced to the mastoids and baseline-corrected using a 100 ms pre-stimulus interval.

A sound was classified as a deviant if it was the first occurrence of a given F0 following a change in the roving paradigm. Five initial sounds in the sequence and every two sounds following a deviant were rejected from the analysis, with the remaining sounds classified as standards. ERPs were computed by averaging responses to standard and to deviant sounds within participants and conditions. Difference waves were derived by subtracting the ERPs of standard sounds from those of deviant sounds for each condition and participant. These difference waves were then used to calculate the amplitudes and latencies for both the MMN and P3a components. Peak measures were calculated on averaged EEG signals from fronto-central electrodes Fz, F1, F2, F3, F4, AF3, AF4, AFz, FC1, FC2 and FCz. MMN latencies were calculated participant- and condition-wise by detecting the latency of the most negative peak (local minimum) in the window from 70 ms to 250 ms post-stimulus. Similarly, P3a latencies were calculated for a positive peak in the window between 150 ms and 350 ms. MMN and P3a amplitudes were calculated as mean amplitudes of the signal between +/− 25 ms around the participant- and condition-wise peak.

We tested whether the MMN and P3 amplitudes differ significantly from zero using one-sample, two-tailed t-tests. *P*-values correction based on the Benjamini-Hochberg procedure was applied to keep the false discovery rate at the 0.05 significance level, separately for the MMN and P3 amplitudes comparisons. To test whether harmonicity affects the amplitude and latency of mismatch responses, we performed a repeated measures ANOVA. Greenhouse–Geisser correction was applied whenever Mauchly’s Test for Sphericity yielded significant results. If ANOVA results showed a significant effect of harmonicity, we performed pairwise post-hoc comparisons using estimated marginal means. P-values were corrected for multiple comparisons using the Tukey method.

To model how increasing inharmonicity affects mismatch responses, we fitted mixed effects models, with differential entropy of the jitter rates distribution, specific to the experimental condition, as the independent variable. Polynomial models up to the fourth degree with random intercepts were fitted for both MMN and P3 amplitude and each model was compared to the lower degree model with a likelihood ratio test. To test the hypothesis that MMN amplitude remains at a similar level for low inharmonicity conditions and then rapidly approaches zero we fitted a mixed effects sigmoid model with random asymptote *L*:

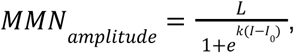

where *I* is the differential entropy of the jitter rates distribution (calculated using log_10_), *I*_0_ is the *I* value at the inflection point, and k determines the slope at the inflection point. First, we fitted a model in which all the parameters were allowed to vary between participants (but were forced to have a diagonal covariance matrix because otherwise the model did not converge). The variances of the k and *I*_0_ parameters were miniscule, so we decided to fit a model with only the asymptote L as a random effect. The values of fixed effects were nearly the same as for the model with random effects for all parameters. To facilitate reproducibility, we report the formulas used for fitting all models in Supplementary Table 6.

All EEG signal processing was performed in Python v3.11, using the packages: MNE v1.5^49,50^, Numpy v1.24^51^, SciPy v1.11^52^ and Pandas v2.1^53^. Data visualization was carried out using Matplotlib v3.7. Modelling and statistical tests were performed in R version 4.3.3 ^54^, with *nlme* ^55^ used for mixed model fitting and *ez* package, version 4.4.0 ^56^ used for ANOVA.

## Author contributions

A.B.: formal analysis, investigation, writing - original draft, writing - review & editing.

B.W.: software, investigation, writing - review & editing.

M.B. - formal analysis, visualisation, writing - original draft, writing - review & editing.

T.D.: investigation, writing - review & editing.

KB: conceptualization, methodology, software, data analysis, data curation, funding acquisition, writing - original draft, review & editing, visualization, supervision.

## Data availability statement

The raw data for this experiment is available at https://doi.org/10.5281/zenodo.15341473 while code that was used to perform the analyses is available at https://doi.org/10.5281/zenodo.17357764.

## Supplementary materials

**Supplementary Table 1.** One-sample, two-sided t-tests of mean MMN and P3 amplitudes for all studied jitter conditions, tested against 0.

| Condition | *M*[95%*CI*] | *t* | *df* | *p* | *p*<sub>adj</sub> |
| --- | --- | --- | --- | --- | --- |
| ***MMN amplitude*** |  |  |  |  |  |
| j0 | -1.17[-1.61, -0.74] | -5.45 | 34 | <.001 | <.001 |
| j1 | -1.49[-1.86, -1.11] | -8.08 | 34 | <.001 | <.001 |
| j2 | -1.13[-1.51, -0.76] | -6.18 | 34 | <.001 | <.001 |
| j3 | -1.22[-1.66, -0.78] | -5.61 | 34 | <.001 | <.001 |
| j4 | -0.91[-1.28, -0.54] | -4.97 | 34 | <.001 | <.001 |
| j5 | -0.37[-0.62, -0.12] | -2.97 | 34 | 0.005 | 0.007 |
| j6 | -0.39[-0.73, -0.06] | -2.39 | 34 | 0.023 | 0.026 |
| j7 | -0.32[-0.64, -0.01] | -2.08 | 34 | 0.045 | 0.045 |
| ***P3 amplitude*** |  |  |  |  |  |
| j0 | 0.93[0.54, 1.32] | 4.83 | 34 | <.001 | <.001 |
| j1 | 0.51[0.12, 0.89] | 2.69 | 34 | 0.011 | 0.011 |
| j2 | 1.01[0.6, 1.42] | 5.00 | 34 | <.001 | <.001 |
| j3 | 0.91[0.55, 1.27] | 5.14 | 34 | <.001 | <.001 |
| j4 | 0.99[0.57, 1.41] | 4.75 | 34 | <.001 | <.001 |
| j5 | 0.79[0.51, 1.07] | 5.71 | 34 | <.001 | <.001 |
| j6 | 0.54[0.19, 0.88] | 3.17 | 34 | 0.003 | 0.004 |
| j7 | 0.58[0.24, 0.92] | 3.48 | 34 | 0.001 | 0.002 |

Note: *M*[95%*CI*] indicates mean and its 95% confidence interval; *t* indicates t-test statistic, *df* indicates degrees of freedom; *p* indicates uncorrected p-value; *p*<sub>adj</sub> was calculated using Benjamini & Hochberg correction for multiple comparisons; results were considered significant if *p*<sub>adj</sub> < 0.05; j0 is the control, harmonic condition and j1-7 indicate subsequent inharmonic conditions.

**Supplementary Table 2.** ANOVA results for the effect of inharmonicity on MMN and P3 amplitudes and latencies.

| Effect | DFn | DFd | SSn | SSd | F | p | ges | W | p<sub>GG</sub> |
| --- | --- | --- | --- | --- | --- | --- | --- | --- | --- |
| ***MMN amplitude*** |  |  |  |  |  |  |  |  |  |
| (Intercept) | 1 | 34 | 214.39 | 121.52 | 59.99 | <.001 | 0.41 |  |  |
| jitter | 7 | 238 | 50.60 | 188.55 | 9.12 | <.001 | 0.14 | 0.17* | <.001 |
| ***P3 amplitude*** |  |  |  |  |  |  |  |  |  |
| (Intercept) | 1 | 34 | 170.46 | 106.25 | 54.54 | <.001 | 0.35 |  |  |
| jitter | 7 | 238 | 10.81 | 206.97 | 1.78 | 0.093 | 0.0 | 0.37 |  |
| ***MMN latency*** |  |  |  |  |  |  |  |  |  |
| (Intercept) | 1 | 34 | 6.85 | 0.05 | 4363.87 | <.001 | 0.94 |  |  |
| jitter | 7 | 238 | 0.01 | 0.35 | 0.82 | 0.57 | 0.02 | 0.2* | 0.538 |
| ***P3 latency*** |  |  |  |  |  |  |  |  |  |
| (Intercept) | 1 | 34 | 18.40 | 0.09 | 7223.28 | <.001 | 0.98 |  |  |
| jitter | 7 | 238 | 0.01 | 0.36 | 1.22 | 0.29 | 0.03 | 0.23* | 0.299 |

Note: DFn - degrees of freedom of the numerator; DFd - degrees of freedom of the denominator; SSn - sum of squares of the numerator; SSd - sum of squares of the denominator; F - F statistic; p - uncorrected p-value; ges - generalized eta squared; W - Mauchly's W statistic; p<sub>GG</sub> - Greenhouse-Geiser corrected p-value.

**Supplementary Table 3.**
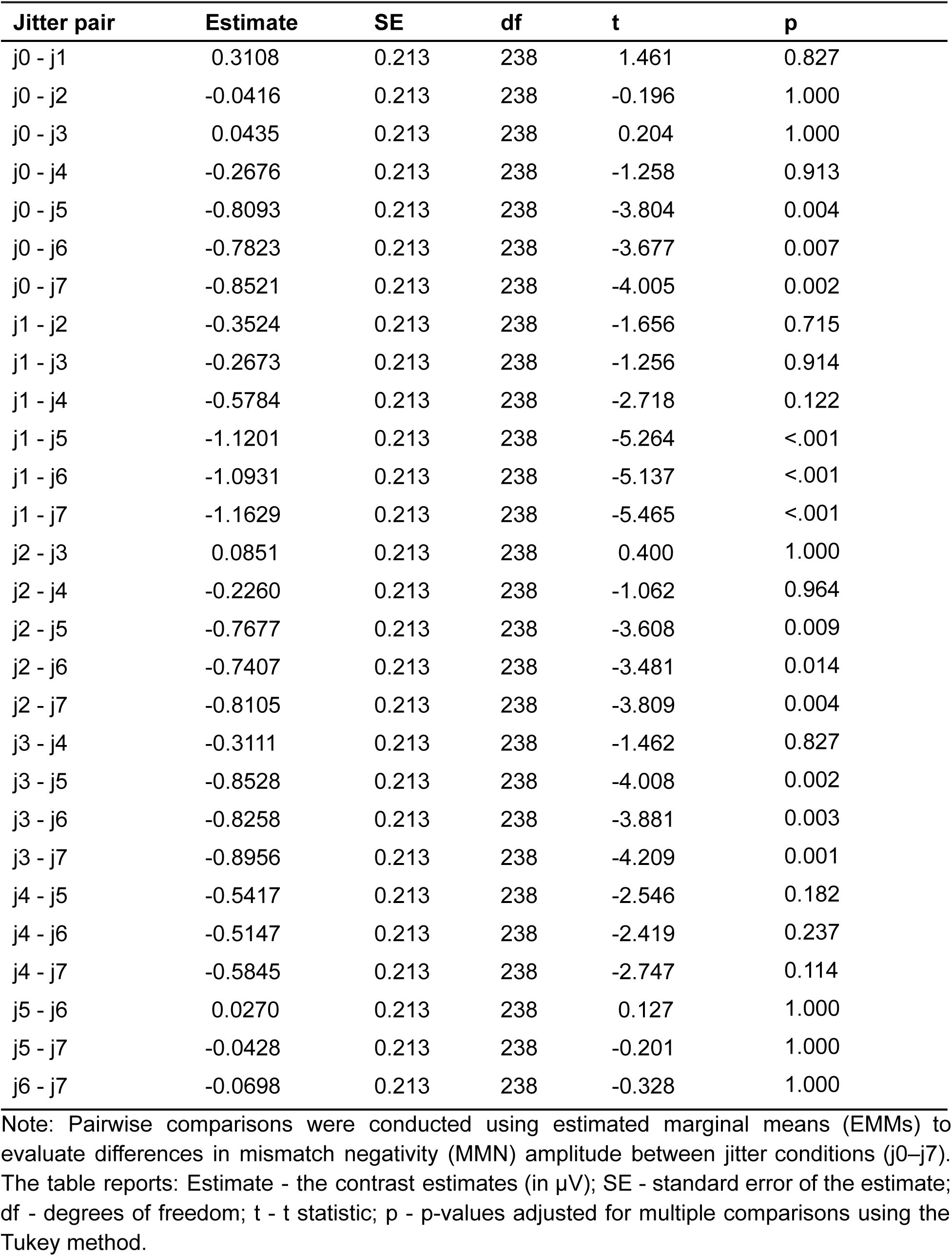
Pairwise contrasts of mismatch negativity amplitude across jitter conditions.

| Jitter pair | Estimate | SE | df | t | p |
| --- | --- | --- | --- | --- | --- |
| j0 - j1 | 0.3108 | 0.213 | 238 | 1.461 | 0.827 |
| j0 - j2 | -0.0416 | 0.213 | 238 | -0.196 | 1.000 |
| j0 - j3 | 0.0435 | 0.213 | 238 | 0.204 | 1.000 |
| j0 - j4 | -0.2676 | 0.213 | 238 | -1.258 | 0.913 |
| j0 - j5 | -0.8093 | 0.213 | 238 | -3.804 | 0.004 |
| j0 - j6 | -0.7823 | 0.213 | 238 | -3.677 | 0.007 |
| j0 - j7 | -0.8521 | 0.213 | 238 | -4.005 | 0.002 |
| j1 - j2 | -0.3524 | 0.213 | 238 | -1.656 | 0.715 |
| j1 - j3 | -0.2673 | 0.213 | 238 | -1.256 | 0.914 |
| j1 - j4 | -0.5784 | 0.213 | 238 | -2.718 | 0.122 |
| j1 - j5 | -1.1201 | 0.213 | 238 | -5.264 | <.001 |
| j1 - j6 | -1.0931 | 0.213 | 238 | -5.137 | <.001 |
| j1 - j7 | -1.1629 | 0.213 | 238 | -5.465 | <.001 |
| j2 - j3 | 0.0851 | 0.213 | 238 | 0.400 | 1.000 |
| j2 - j4 | -0.2260 | 0.213 | 238 | -1.062 | 0.964 |
| j2 - j5 | -0.7677 | 0.213 | 238 | -3.608 | 0.009 |
| j2 - j6 | -0.7407 | 0.213 | 238 | -3.481 | 0.014 |
| j2 - j7 | -0.8105 | 0.213 | 238 | -3.809 | 0.004 |
| j3 - j4 | -0.3111 | 0.213 | 238 | -1.462 | 0.827 |
| j3 - j5 | -0.8528 | 0.213 | 238 | -4.008 | 0.002 |
| j3 - j6 | -0.8258 | 0.213 | 238 | -3.881 | 0.003 |
| j3 - j7 | -0.8956 | 0.213 | 238 | -4.209 | 0.001 |
| j4 - j5 | -0.5417 | 0.213 | 238 | -2.546 | 0.182 |
| j4 - j6 | -0.5147 | 0.213 | 238 | -2.419 | 0.237 |
| j4 - j7 | -0.5845 | 0.213 | 238 | -2.747 | 0.114 |
| j5 - j6 | 0.0270 | 0.213 | 238 | 0.127 | 1.000 |
| j5 - j7 | -0.0428 | 0.213 | 238 | -0.201 | 1.000 |
| j6 - j7 | -0.0698 | 0.213 | 238 | -0.328 | 1.000 |

Note: Pairwise comparisons were conducted using estimated marginal means (EMMs) to evaluate differences in mismatch negativity (MMN) amplitude between jitter conditions (j0–j7). The table reports: Estimate - the contrast estimates (in $\mu\text{V}$ ); SE - standard error of the estimate;
df - degrees of freedom; t - t statistic; p - p-values adjusted for multiple comparisons using the Tukey method.

**Supplementary Table 4.** Model comparison of MMN and P3 amplitude as a function of jitter.

| Model degree | df | AIC | BIC | logLik | Test | L.Ratio | p |
| --- | --- | --- | --- | --- | --- | --- | --- |
| ***MMN amplitude - linear and polynomial models*** |  |  |  |  |  |  |  |
| 0 (Intercept only) | 3 | 736.13 | 746.63 | -365.07 |  | - | - |
| 1 (linear) | 4 | 691.23 | 705.24 | -341.62 | 0 vs 1 | 46.90 | <.001 |
| 2 (quadratic) | 5 | 693.01 | 710.51 | -341.50 | 1 vs 2 | 0.23 | 0.635 |
| 3 (cubic) | 6 | 693.66 | 714.67 | -340.83 | 2 vs 3 | 1.35 | 0.246 |
| 4 (quartic) | 7 | 694.77 | 719.28 | -340.38 | 3 vs 4 | 0.89 | 0.345 |
| ***MMN amplitude - sigmoid model*** |  |  |  |  |  |  |  |
| 0 (Intercept only) | 3 | 736.13 | 746.63 | -365.07 |  | - | - |
| Sigmoid model | 5 | 664.65 | 682.15 | -327.32 | 0 vs 1 | 75.48 | <.001 |
| ***P3 amplitude - linear and polynomial models*** |  |  |  |  |  |  |  |
| 0 (Intercept only) | 3 | 716.43 | 726.93 | -355.22 |  | - | - |
| 1 (linear) | 4 | 717.46 | 731.46 | -354.73 | 0 vs 1 | 0.97 | 0.324 |
| 2 (quadratic) | 5 | 713.50 | 731.01 | -351.75 | 1 vs 2 | 5.95 | 0.015 |
| 3 (cubic) | 6 | 712.62 | 733.63 | -350.31 | 2 vs 3 | 2.88 | 0.090 |
| 4 (quartic) | 7 | 714.62 | 739.13 | -350.31 | 3 vs 4 | 0.00 | 0.966 |

Note: df - degrees of freedom; AIC - Akaike information criterion; BIC - bayesian information criterion; logLik - log-likelihood; Test indicates which models (degrees) are compared in the test; L.Ratio - likelihood ratio; p - p-value of the likelihood ratio test.

**Supplementary Table 5.**
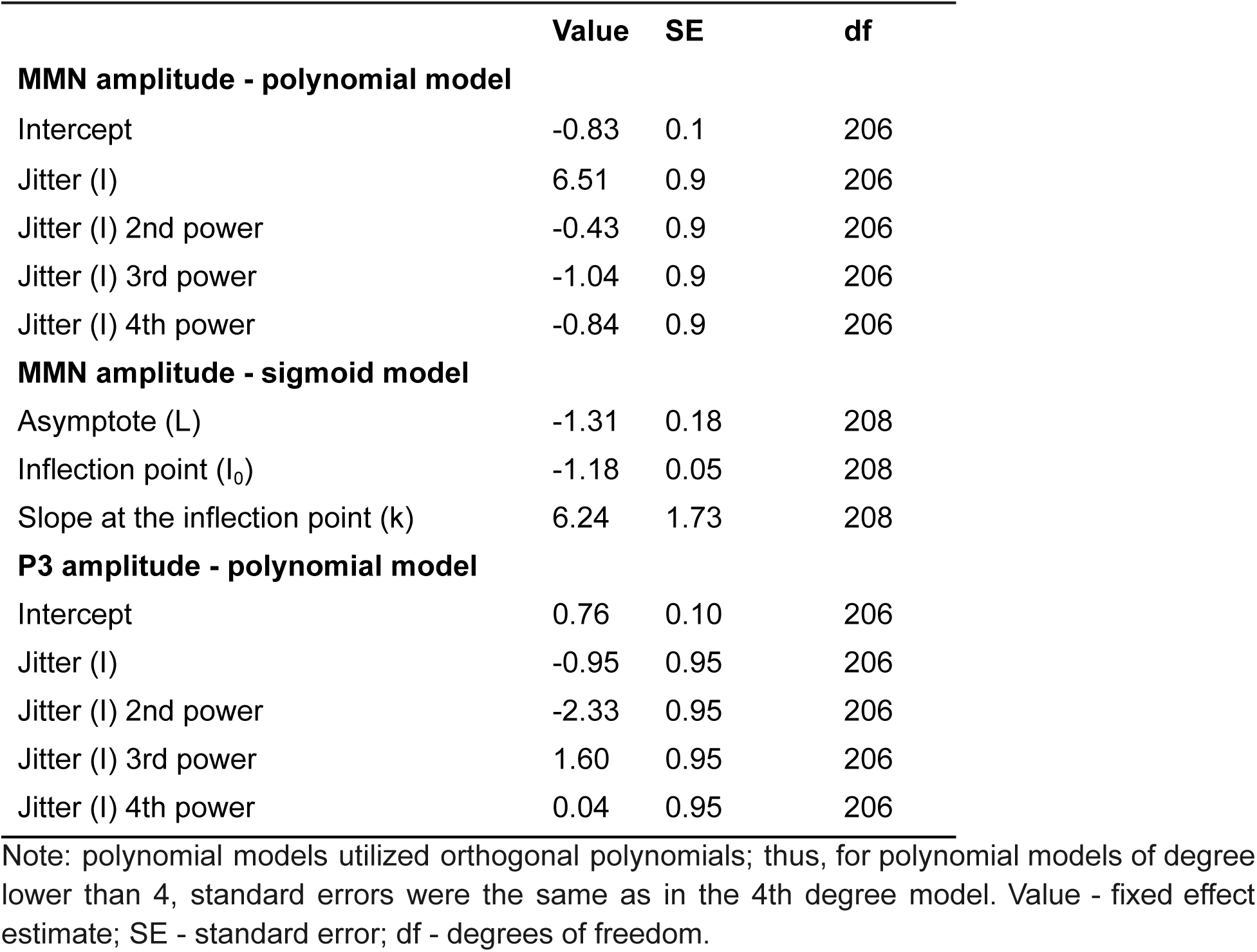
Fitted models fixed effects.

**Supplementary Table 6.** R formulas used for fitting linear, polynomial and sigmoid models.

| **Model degree** | **Formula** |
| --- | --- |
| ***MMN amplitude - linear and polynomial models*** |  |
| 0 (Intercept only) | mmn_amp ~ 1 |
| 1 (linear) | mmn_amp ~ poly(jitter, 1) |
| 2 (quadratic) | mmn_amp ~ poly(jitter, 2) |
| 3 (cubic) | mmn_amp ~ poly(jitter, 3) |
| 4 (quartic) | mmn_amp ~ poly(jitter, 4) |
| ***MMN amplitude - sigmoid model*** |  |
| 0 (Intercept only) | mmn_amp ~ 1 |
| Sigmoid model | mmn_amp ~ sigmoid(jitter, L, I_0, k) |
| ***P3 amplitude - linear and polynomial models*** |  |
| 0 (Intercept only) | p3_amp ~ 1 |
| 1 (linear) | p3_amp ~ poly(jitter, 1) |
| 2 (quadratic) | p3_amp ~ poly(jitter, 2) |
| 3 (cubic) | p3_amp ~ poly(jitter, 3) |
| 4 (quartic) | p3_amp ~ poly(jitter, 4) |

Note: For all models, participant id was inserted as random effect.

**Supplementary Figure 1.**
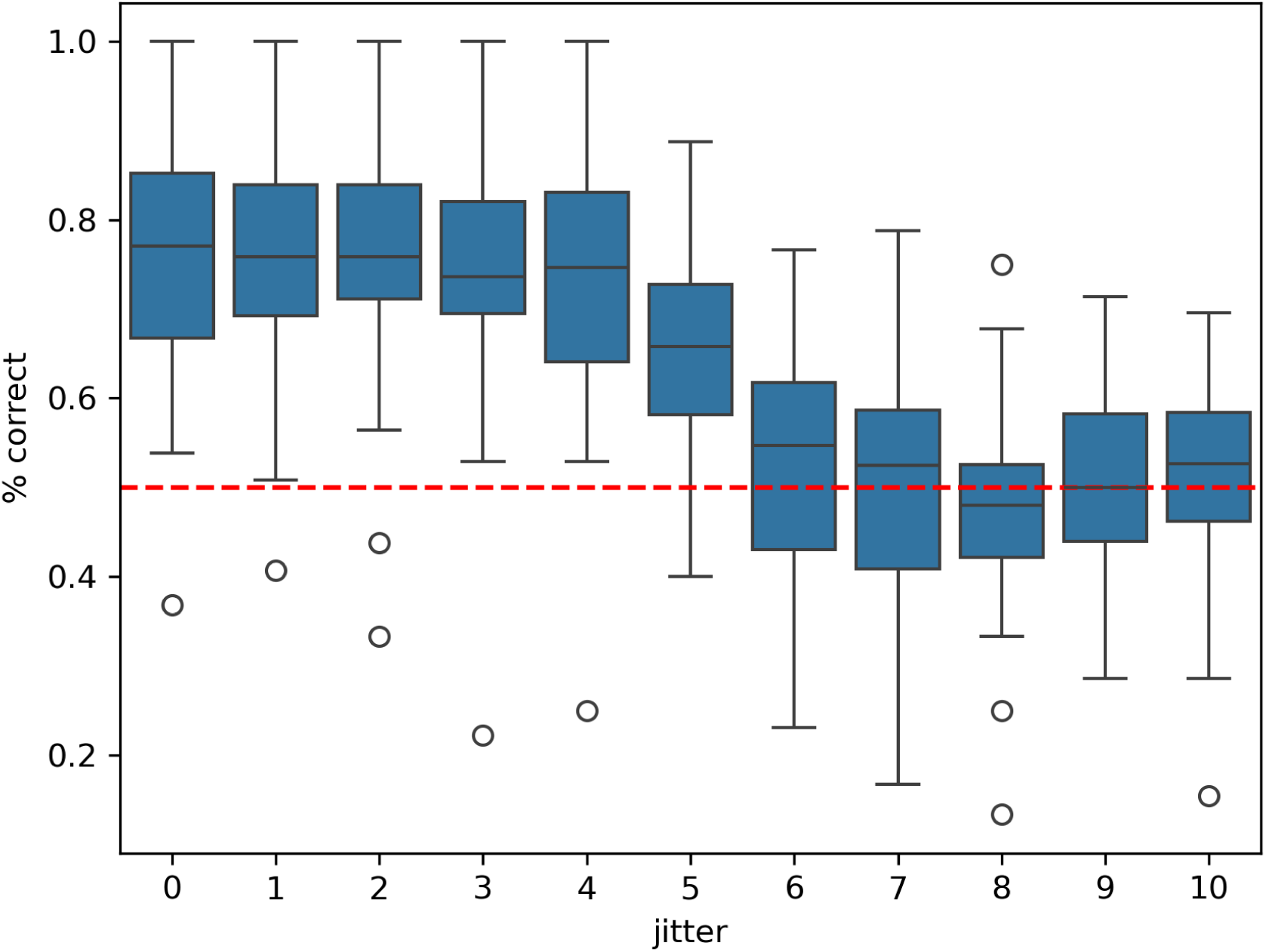
Results of the behavioral experiment. Figure shows percent-correct responses as a function of jitter. Boxes represent the range of quartiles 2 and 3, horizontal line within the box represents the median while whiskers indicate minima and maxima excluding outliers (shown as circles). Red dashed line indicates chance-level performance (50%).

**Supplementary Figure 2.**
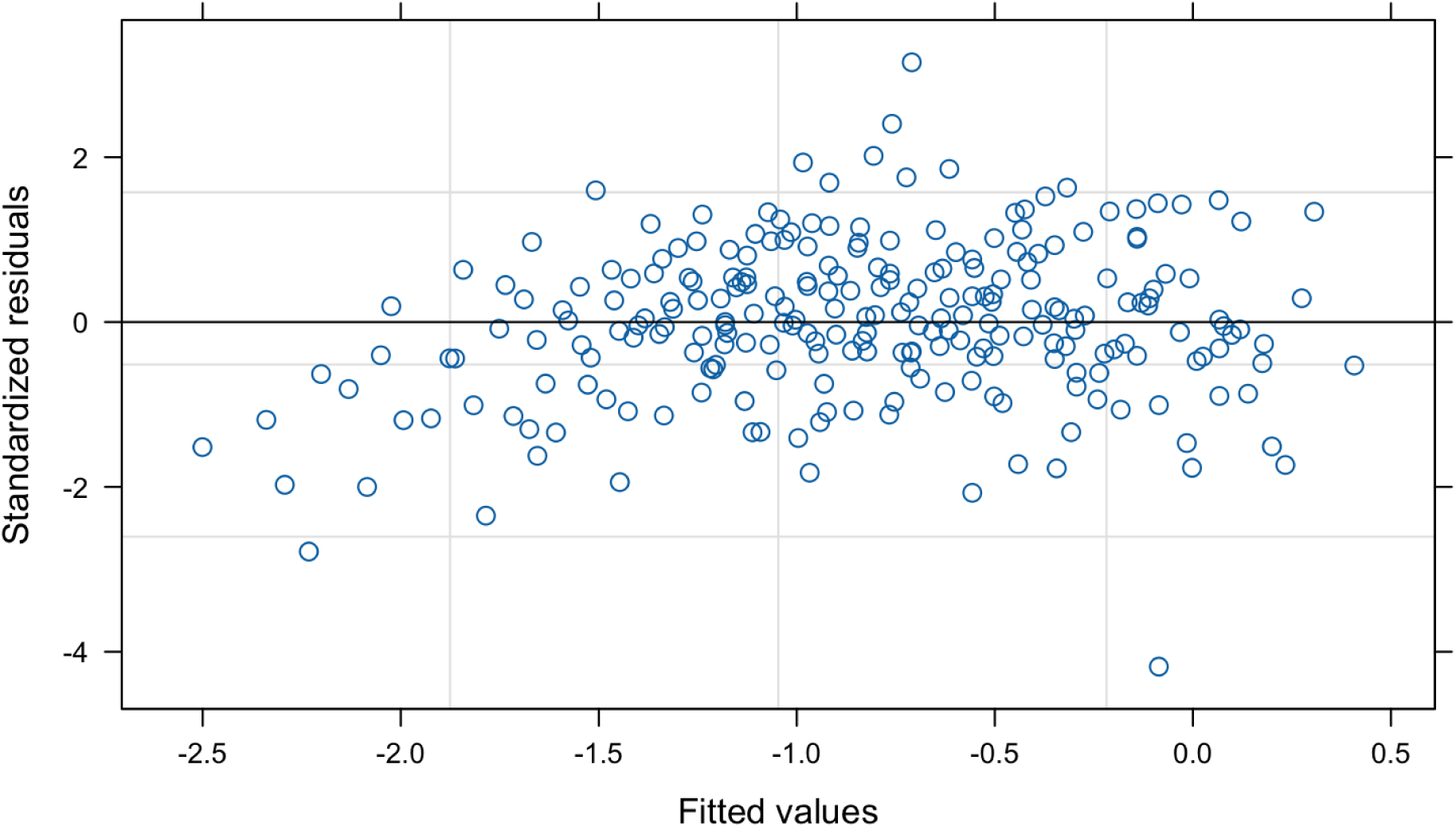
Residual plot for the linear model of MMN amplitude. Fitted values are presented on x-axis while standardized residuals are presented on y-axis.

**Supplementary Figure 3.**
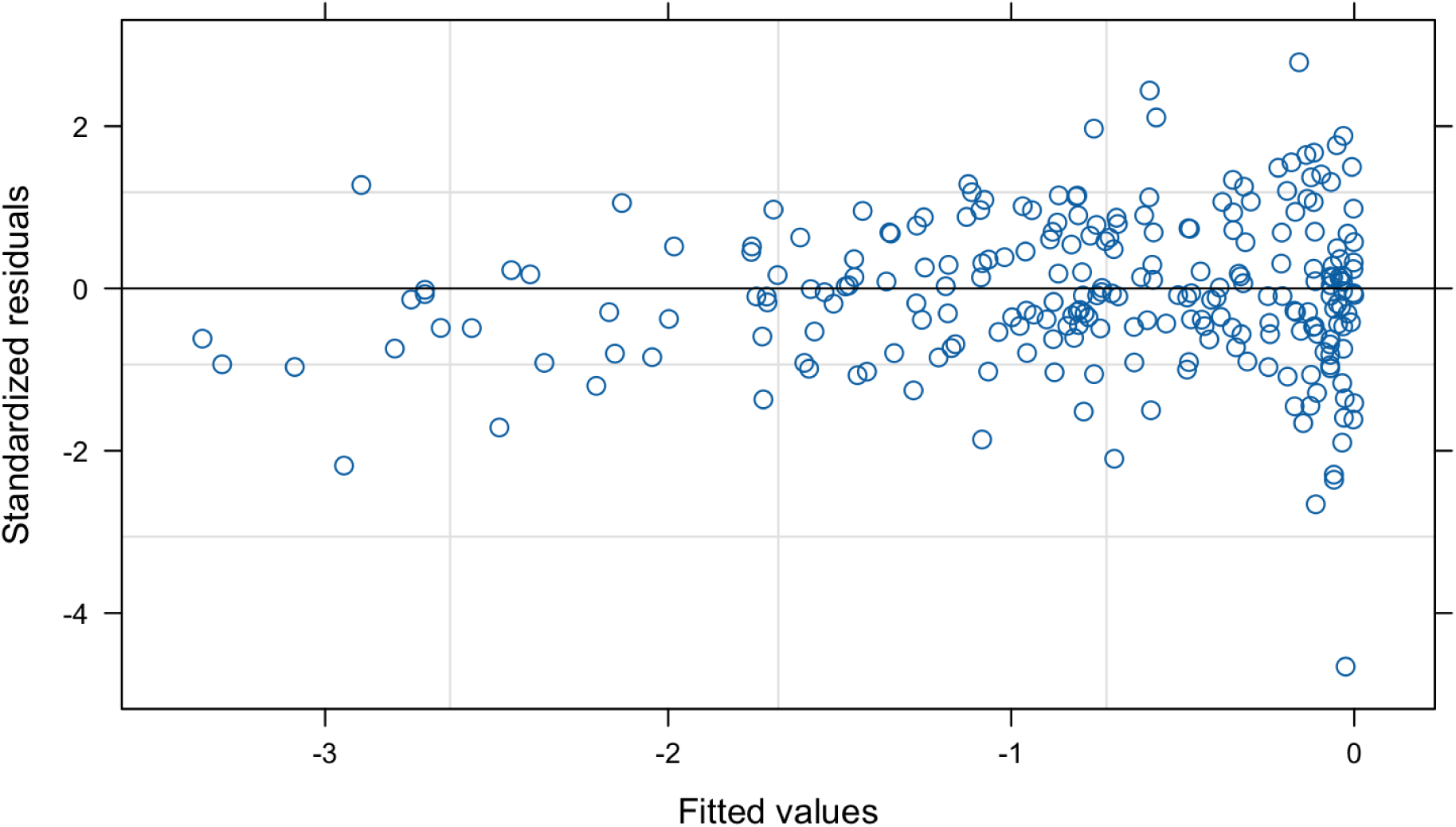
Residual plot for the sigmoid model of MMN amplitude. Fitted values are presented on x-axis while standardized residuals are presented on y-axis.

